# N6-Adenosine Methylation of a Single Site in SARS-CoV-2 5′-UTR Promotes Translation

**DOI:** 10.1101/2022.10.17.512569

**Authors:** Ammar Aly, Gary Scott, Mario Calderon, A. Pejmun Haghighi

## Abstract

The coronavirus disease 2019 (COVID19) led to devastating health outcomes and has continued to spread despite global vaccination efforts^1^. This, alongside the rapid emergence of vaccine resistant variants, creates a need for orthogonal therapeutic strategies targeting more conserved facets of severe acute respiratory syndrome coronavirus (SARS-CoV-2)^2–7^. The viral genome is a single positive RNA strand divided into a genomic and a subgenomic segment. All 16 non-structural viral proteins are translated from the genomic polycistronic ORFs 1a, and 1b using a single 5′-UTR leader^8,9^. To our surprise, the full length 5′-UTR efficiently initiates protein translation despite its predicted structural complexity. Through a combination of biochemical assays and bioinformatic analyses, we demonstrate that a single METTL3-dependent m^6^A methylation event in SARS-CoV-2 5′-UTR regulates the rate of translation initiation. We demonstrate that m^6^A likely exerts this effect by destabilizing the third stem loop (SL3) and increasing overall accessibility to protein complexes. Our discovery provides a foundational insight into the biology of SARS-CoV-2 by linking m^6^A modification to its translational regulation and, thus, opens a new avenue for potential novel therapeutic strategies.

## INTRODUCTION

COVID-19 continues to exert an extraordinary global impact, with nearly 780 million confirmed cases and more than 7 million reported deaths to date^10^. Despite unprecedented efforts to vaccinate populations worldwide, the viral spread persists—primarily driven by the emergence and spread of variants capable of evading vaccine-induced immunity^11,12^. This troubling trend reflects a confluence of factors: uneven vaccine distribution and uptake, relaxation of non-pharmaceutical interventions (such as mask mandates and physical distancing), and high rates of viral transmission in partially immunized communities. Under these conditions, SARS-CoV-2 variants that harbor mutations conferring both increased transmissibility and immune escape gain a selective advantage^12^.

This evolutionary pressure is most pronounced in the viral spike (S) glycoprotein—the principal target of all vaccines authorized in the United States and European Union. The spike protein’s receptor-binding domain and adjacent epitopes are inherently hypermutable^6,11–13^, permitting the rapid accrual of substitutions that diminish neutralizing antibody binding. Indeed, successive variants of concern—Delta, Omicron, XBB lineage, and most recently JN.1—have each accrued distinct constellations of spike mutations that enhance ACE2 receptor affinity and reduce vaccine-elicited antibody recognition, thereby increasing both infectivity and virulence^11,12^.

In light of these challenges, combinatorial antiviral strategies have emerged as a promising approach to suppress the evolution of treatment-resistant strains. By applying multiple, independent selective pressures—analogous to the multidrug regimens that transformed the management of HIV^14^, HBV^15^, tuberculosis^16^, and certain cancers^17^—such therapies elevate the mutational barrier required for viral escape. For SARS-CoV-2, identifying viral targets that are both functionally indispensable and evolutionarily constrained is therefore a critical priority to complement existing vaccination campaigns.

One such conserved feature is the mechanism of discontinuous transcription, a hallmark of coronavirus genome replication^18^. This process hinges on a short Transcription Regulatory Sequence Leader (TRS-L) in the 5′-untranslated region (UTR) of the viral genome, which base-pairs with complementary TRS-body (TRS-B) sequences upstream of each open reading frame (ORF)^8,9,19–21^. During negative-strand synthesis, the replicase complex switches templates from TRS-B sites to the TRS-L, generating a nested set of Subgenomic RNAs (sgRNAs) that are subsequently transcribed into individual viral mRNAs. Because every sgRNA—spanning structural and accessory genes—requires the same TRS-L segment for its synthesis, the 5′-UTR remains highly conserved across the Coronaviridae family, both in primary nucleotide sequence and in predicted secondary structure^8,18,21–23^. Moreover, the non-structural viral proteins are translated from the polycistronic ORFs 1a, and 1b, which utilize the same 265 bp region as their 5′-UTR. The remaining transcripts also known as sub-genomic transcripts contain 5′-UTRs of varying lengths with the 5′ 1-75 nucleotides common to all^9^. Indeed, sequence alignments reveal minimal variation in the 5′-UTR among early arising SARS-CoV-2 variants (Supplementary Figure 1).

Despite the central role of 5′-UTRs in regulating mRNA translation^24–26^, we know little about the regulatory influence of SARS-CoV-2 5′-UTR on translation. Secondary structure analyses^21,23,27–29^ predict a complex architecture of stem-loops and pseudoknots in the full-length 5′-UTR. Such complexity would ordinarily impede translation initiation, which relies on 5′-to-3′ scanning by the pre-initiation complex and ATP-dependent unwinding by the eIF4A helicase^30–32^. In cellular mRNAs, extensive 5′-UTR structure correlates strongly with reduced translational efficiency^33^; however, coronaviral mRNAs paradoxically achieve high translation rates and outcompete host transcripts for ribosomal engagement^22,34–36^.

We reasoned that specific RNA modifications might underlie this apparent contradiction. N^6^-methyladenosine (m^6^A) is the most prevalent internal modification in eukaryotic and viral mRNAs^37^, comprising approximately 99% of all known modifications. First detected in viral RNAs in 1974^38^, m^6^A has only recently been linked to the life cycles of diverse viruses^39–44^. The “writer” complex, centered on METTL3, deposits m^6^A at consensus DRACH motifs ([A/G][A/G]AC[U/A/C]) in a context-dependent manner^37,45,45^ after which “reader” proteins (YTHDF1–3) interpret these marks to influence RNA splicing, stability, localization, and translation^46^. Recent studies have mapped m^6^A sites in host transcriptomes during SARS-CoV-2 infection^47^, as well as in the viral genome itself^48^, but few have focused specifically on the 5′-UTR^49^.

To fill this gap, we engineered a series of SARS-CoV-2 5′-UTR variants with targeted alterations at predicted m^6^A sites and assessed their impact on translation using luciferase reporter assays. Our data confirm that, despite its predicted intricate secondary structure, the SARS-CoV-2 5′-UTR is capable of driving robust translation. Strikingly, mutation of a single m^6^A site within the 5′-UTR dramatically reduces translational output, an outcome mimicked by pharmacological inhibition of METTL3. Contrary to our expectation, this effect is not mediated by YTHDF reader proteins; rather, m^6^A appears to facilitate translation likely by destabilizing local RNA structure. Based on these findings, we propose a model in which m^6^A modification of the SARS-CoV-2 5′-UTR fine-tunes viral translational efficiency, thereby highlighting the viral 5′-UTR as a potential target for future COVID-19 therapeutics.

## RESULTS

### SARS-CoV-2 5′-UTR is unexpectedly efficient at driving protein translation

Computational RNA modeling algorithms predict a highly complex secondary structure for the SARS-CoV-2 5′-UTR. To place it in the broader landscape of 5′-UTR structural diversity, we compared it to a 4-fold concatemer of the human ß-globin (hbb) 5′-UTR (Figure 1A), which is an efficient promoter of translation, that has also served as the canonical leader sequence in many of the current mRNA vaccine platforms^50–52^. We also employed genome-wide secondary structure prediction across all annotated human 5′-UTRs, computing a length-normalized stability metric (MFE per nucleotide) to correct for the intrinsic bias of longer sequences toward more negative free energy values^53^. Focusing on the subset of human transcripts whose GC content (40–50 %) and length (150–275 nt) closely resemble those of the SARS-CoV-2 5′-UTR (GC = 44.5 %, length = 265 nt) and 4-fold Hbb 5′-UTR (GC = 44 %, length = 200 nt), we observed that the SARS-CoV-2 5′-UTR resides in the upper tail of the distribution for structural density (FDR < 0.05; Figure 1B). This indicates that, on a per-base basis, it forms a more densely packed network of secondary structures than over 95% of human transcripts of comparable composition.

**Figure 1.**
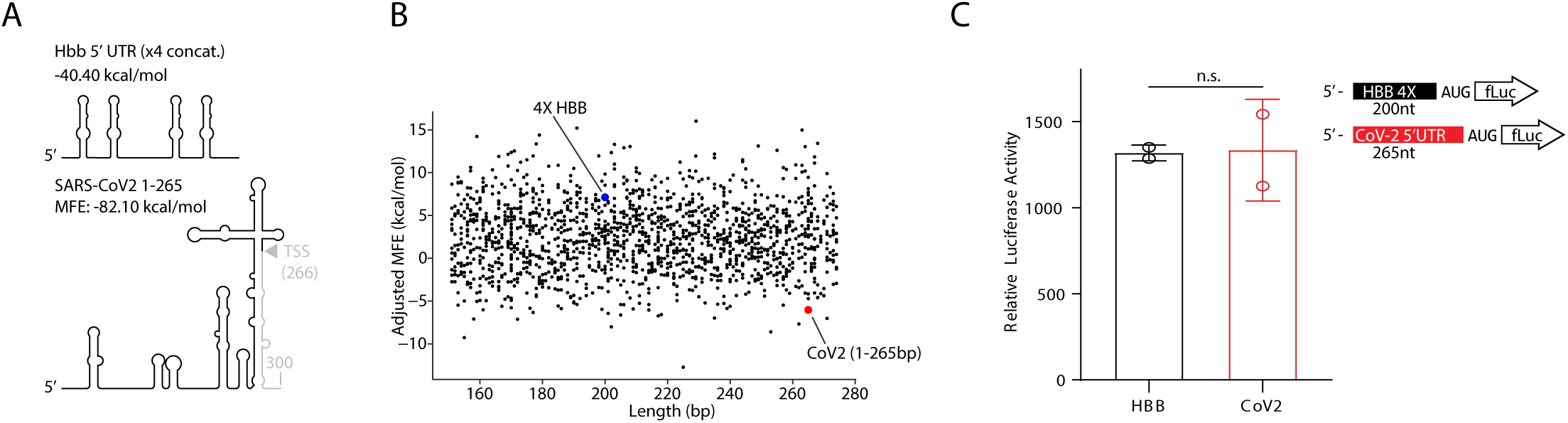
SARS-CoV-2 5’-UTR Is an Unexpectedly Efficient Promoter of Translation. Secondary structure and minimum free energy of **A.** 4X repeat of Hbb 5’-UTR and SARS-CoV-2 5’-UTR (bp 1-265).**B.** Adjusted Minimum Free Energy for human 5’-UTRs with GC Content of ∼50% and Length between 150-280bp. Hbb (62% GC) and CoV-2 (47% GC) marked in blue and red respectively. **C.** Luciferase assay from pGL3-Hbb-fLuc and pGL3-CoV-2 5’-UTR-fLuc transfected HEK293T cells. Results of two experiments (3 biological replicates each) shown as mean±SEM. Statistical testing was performed using Welch’s T-test.

Given the well-established inverse relationship between 5′-UTR secondary structure complexity and translational efficiency—whereby those with more stable secondary structure and lower minimum free energy (MFE) typically impede ribosomal scanning and initiation—we predicted that the SARS-CoV-2 5′-UTR (MFE of –82.10 kcal/mol) would drive lower protein output than the relatively unstructured 4X Hbb 5′-UTR (MFE of –40.40 kcal/mol) leader^33^. To test this hypothesis, we cloned each 5′-UTR directly upstream of a firefly luciferase open reading frame in a standardized expression vector and transfected these constructs into HEK293T cells. Quantitative luminescence assays performed 24 hours post-transfection revealed, to our surprise, no statistically significant difference in luciferase activity or mRNA levels between the SARS-CoV-2 and 4X Hbb 5′-UTR reporters (Figure 1C and Supplementary Figure 2A), suggesting that the viral leader harbors additional features—beyond its predicted thermodynamic stability—that enable robust translation in a cellular context.

### SARS-CoV-2 5′-UTR m^6^A methylation enhances protein translation efficiency

Given the abundance of m^6^A modification in eukaryotic and viral transcripts^37^and its destabilizing effect in regions of RNA that are base-paired^54,55^, we reasoned that m^6^A methylation might underlie the unexpectedly high translational efficiency of the SARS-CoV-2 5′-UTR. To test this hypothesis, we first asked whether the viral leader is decorated with m^6^A marks. In silico scans for the consensus DRACH motif across the 265-nt 5′-UTR, coupled with established m^6^A site predictors, revealed a single high-confidence candidate at adenine 74, embedded within the essential TRS-L sequence in a base-paired region (Figure 2c)^56^. Next, we performed m^6^A RNA immunoprecipitation (MeRIP) on total RNA harvested from HEK293T cells transfected with a SARS-CoV-2 5′-UTR–firefly luciferase reporter. Quantitative RT-PCR of the anti-m^6^A pulldown fraction showed a robust ∼52-fold enrichment of the viral leader relative to input (mean ± SEM: 52.0 ± 10.2; p < 0.001), confirming that the 5′-UTR is indeed methylated in these cells (Figure 2A, B).

**Figure 2.**
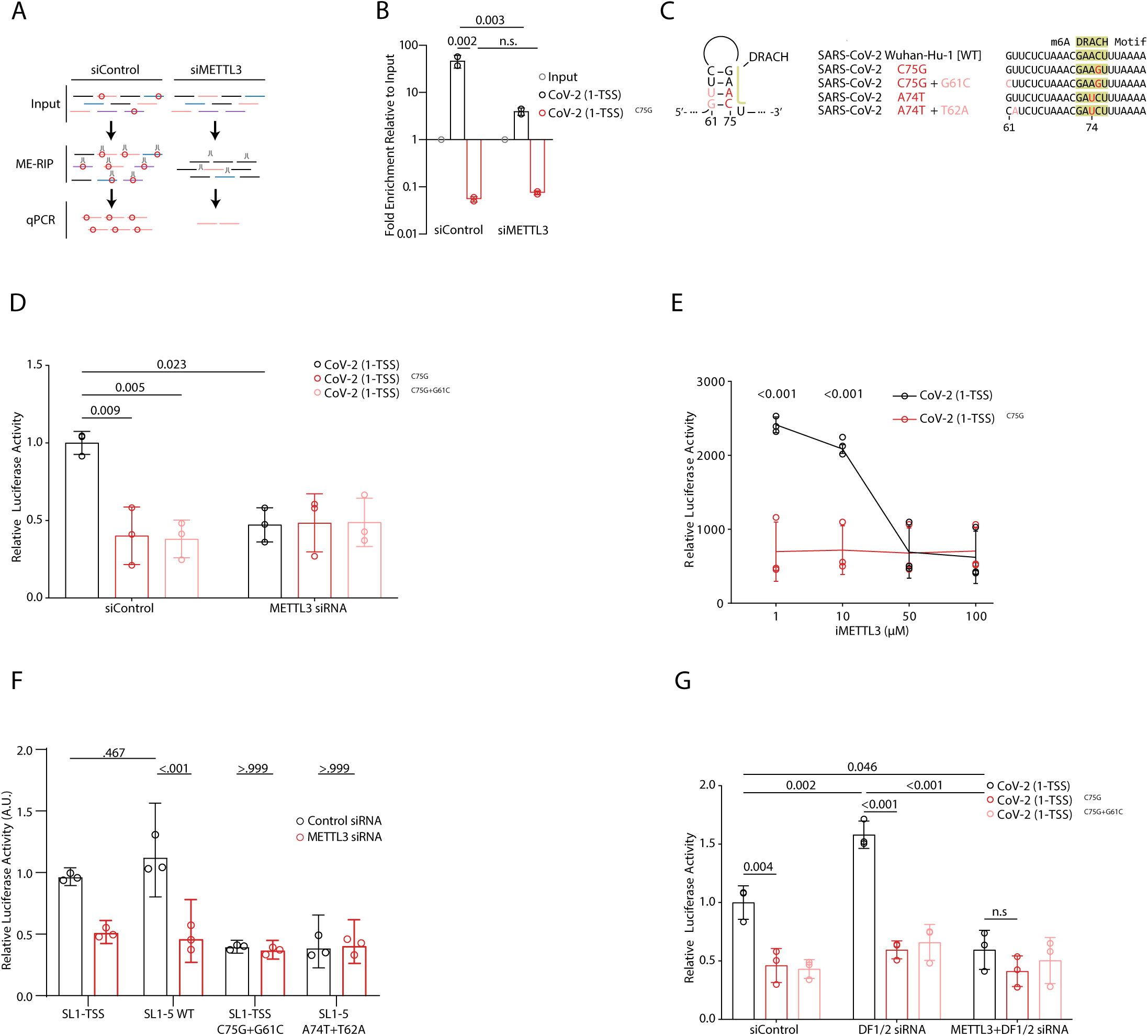
m^6^A enhances SARS-CoV-2 5’-UTR Initiated Translation. **A.** Experimental design for m^6^A-RNA Immunoprecipitation (ME-RIP) and RT-qPCR for SARS-CoV-2 5’-UTR. **B.** ME-RIP-RT-qPCR enrichment of CoV-2 5’-UTR from HEK293T cells transfected with pGL3-CoV-2-5’-UTR (SL1-TSS)-fLuc or pGL3-CoV-2-5’-UTR^C75G^ (SL1-TSS)-fLuc treated with and without METTL3 siRNA knockdown. Results of two experiments (3 biological replicates each) normalized to input fractions shown as geometric mean; error bars are geometric SD. Statistical testing was performed using Tukey’s HSD under two-way ANOVA. **C.** Sequences of putative m^6^A site in SARS-CoV2 WT and engineered m^6^A mutant variants and predicted DG for respective 5’UTRs (1-265). **D.** Luciferase assay of pGL3-CoV-2-5’-UTR (SL1-TSS)-fLuc or pGL3-CoV-2-5’-UTR^C75G^ (SL1-TSS)-fLuc or pGL3-CoV-2-5’-UTR^C75G+G61C^ (SL1-TSS)-fLuc transfected HEK293T with or without METTL3 siRNA knockdown. Results from three experiments (3 biological replicates each) shown as mean ± SEM. Statistical testing was performed using Tukey’s HSD under two-way ANOVA **E.** Luciferase assay of pGL3-CoV-2-5’-UTR (SL1-TSS)-fLuc or pGL3-CoV-2-5’-UTR^C75G^ (SL1-TSS)-fLuc HEK293T treated with increasing concentrations of METTL3 Inhibitor STM2457. Results from three experiments (3 biological replicates each) shown as geometric mean; error bars are geometric SD. Statistical testing was performed using Bonferroni-Šidák correction under two-way ANOVA **F.** Luciferase assay of pGL3-CoV-2-5’-UTR (SL1-TSS)-fLuc, pGL3-CoV-2-5’-UTR^C75G+G61C^ (SL1-TSS)-fLuc, pGL3-CoV-2-5’-UTR (SL1-5)-fLuc, and pGL3-CoV-2-5’-UTR^C75G+G61C^ (SL1-5) -fLuc. Results from three experiments (3 biological replicates each) shown as geometric mean; error bars are geometric SD. Statistical testing was performed using Tukey’s HSD under two-way ANOVA. **G.** Luciferase assay of pGL3-CoV-2-5’-UTR (SL1-TSS)-fLuc or pGL3-CoV-25’-UTR^C75G^ (SL1-TSS)-fLuc or pGL3-CoV-2-5’-UTR^C75G+G61C^ (SL1-TSS)-fLuc transfected HEK293T with or without METTL3 and/or DF1/2 siRNA knockdown. Results from three experiments (3 biological replicates each) shown as geometric mean; error bars are geometric SD. Statistical testing was performed using Tukey’s HSD under two-way ANOVA.

To demonstrate that this enrichment reflects specific methylation at A74, we introduced a point mutation (C75G) that disrupts the DRACH consensus while preserving the adjacent sequence context (Figure 2C). Strikingly, the C75G mutant abolished MeRIP enrichment almost entirely (0.061 ± 0.005; WT vs. C75G p = 0.002), indicating that methylation at A74 is responsible for the observed signal. Consistent with this, siRNA-mediated knockdown of METTL3 reduced enrichment of the WT 5′-UTR from 52.04 ± 10.2 to 4.40 ± 0.49 (p = 0.003), whereas the C75G mutant remained unenriched regardless of METTL3 levels (Figure 2B). These results establish that A74 is a bona fide METTL3-dependent m⁶A site.

Having confirmed m⁶A modification at A74, we next assessed its functional impact on translation. HEK293T cells were transfected with luciferase reporter constructs bearing either the WT 5′-UTR or the C75G mutant. Luciferase assays revealed a dramatic ∼60% reduction in translation for the C75G mutant relative to WT (WT = 1.00 ± 0.043 RLU; C75G = 0.401 ± 0.107 RLU; p = 0.009; Figure 2D). To rule out the possibility that this effect arose from disruption of a predicted base-pair (G61–C75), we generated a compensatory double mutant (C75G + G61C) that restores the pairing but eliminates the DRACH motif. This double mutant exhibited an identical translational defect to C75G alone, confirming that loss of m^6^A, and not secondary-structure perturbation, underlies the diminished luciferase output (Figure 2D). We found no significant differences in mRNAs between the different conditions suggesting that translational, and not transcriptional, effects are responsible for the different luciferase levels (Supplementary Figure 2B).

In support of the critical role of m^6^A modification in translational regulation, we found that METTL3 siRNA knockdown phenocopied the effect of elimination of the DRACH motif by suppressing the ability of the WT SARS-CoV2 5′-UTR to drive translation (siControl = 1.00 ± 0.043 vs. siMETTL3 = 0.47 ± 0.064 RLU; p = 0.023) without causing further suppression of the C75G mutant 5′-UTR (Figure 2D). Similarly, pharmacological inhibition of METTL3 with the specific small-molecule STM2457 produced a dose-dependent suppression of WT 5′-UTR-driven translation, while leaving the mutant 5′-UTR^C75G^-driven translation unchanged (Figure 2E). We found no change in luciferase reporter mRNA levels between any of the conditions, further indicating the observed differences in luciferase activity were a result of translational regulation (Supplementary Figure 2C). The lack of additive effect of DRACH mutation and pharmacological inhibition supports the conclusion that m^6^A modification at A74, and not at other potential methylation sites, accounts for the translational enhancement conferred by the viral leader. Furthermore, these results rule out any significant contribution from the effect of METTL3 knockdown on the host machinery.

The SARS-CoV-2 5′-UTR as typically defined spans nucleotides 1–265 (SL1-TSS) of the viral genome, terminating at the translational_start site (TSS) of ORF1. However, structural and biochemical probing studies have demonstrated that an additional 35-nt segment immediately downstream of the TSS participates in the formation of the fifth stem–loop (SL5), completing the native in vivo secondary structure of the UTR sequence^21,57–59^. To determine whether inclusion of this SL5 extension alters methylation-dependent regulatory features, we cloned an “extended” 5′-UTR (nt 1–300) upstream of the luciferase reporter (SL1-SL5) and performed the same suite of translation assays. The SL1-SL5 leader recapitulated the translational regulation we observed with 1-265 leader (Figure 1C) and allowed for efficient translation of luciferase (Figure 2F). To test the role of m^6^A modification at A74, we mutated this site and the complimentary nucleotide on the opposite strand to disrupt the DRACH motif and maintain the integrity of the stem loop (SL1-SL5^A74T+T62A^). The 5′-UTR^A74T+T62A^ mutant behaved similarly to the 5′-UTR^C75G+G61C^ mutant, showing impaired translation efficiency compared to the control SL1-SL5 leader and remained unresponsive to METTL3 knockdown (Figure 2F). Both the SL1-TSS and SL1-5 constructs produced similar levels of mRNA, ruling out the possibility that an unexpected transcriptional difference underlies our observations (Supplementary Figure 2D). Together, these data confirm that the canonical 265 nt 5′-UTR and C75G mutant faithfully represent the translational regulation of the extended SARS-CoV-2 5′-UTR—including the complete SL5—and loss of A74 methylation. For the remainder of the experiments, we used these two leader constructs interchangeably.

Finally, because canonical m^6^A biology invokes YTH-domain “reader” proteins to mediate downstream effects, we tested whether two prominent cytoplasmic readers YTHDF1 and YTHDF2 are required for 5′-UTR activity. Surprisingly, siRNA knockdown of both readers in HEK293T cells led to a ∼1.6-fold increase in luciferase translation from the WT 5′-UTR (siControl = 1.00 ± 0.083 vs. DF1/2 siRNA = 1.58 ± 0.067 RLU; p = 0.002; Figure 2G), an effect that was lost when m^6^A was abrogated by either C75G mutation or METTL3 knockdown. We found no change in mRNA levels for any of the constructs following either METTL3 or DF1/DF2 knockdown (Supplementary Figure 2E). Our findings indicate that rather than acting as positive effectors of METTL3, YTHDF proteins appear to antagonize 5′-UTR-mediated translation in this context. This reveals an antagonistic interplay between m^6^A modification, which promotes translation, and reader engagement at A74, which appears to hinder translation. Furthermore, these results suggest that mere methylation of A74 by Mettle3 is sufficient to promote translation independently of YTHDF readers. While the action of the readers appears to be dependent on m^6^A modification at A74, we cannot at this point determine at which point in the process of translation they exert their inhibitory influence. We did not pursue the role of readers in this regulation any further in the current work.

### Loss of SARS-CoV-2 5′-UTR methylation negatively affects association with polysomes

The number of ribosomes associated with a given transcript is a reliable measure of translational efficiency of that transcript; this is estimated by determining the position of the peak of gradient fractions containing that transcript in a polysome fractionation experiment. To build on our discovery that m^6^A enhances SARS-CoV-2 5′-UTR–mediated translation, we performed ribosome profiling to map the changes in ribosome association of our reporter mRNAs in HEK293T cells under conditions of low and high m^6^A modification.

Polysome profiles generated from total HEK293T cell extracts showed indistinguishable patterns of 40S, 60S, 80S, and polysome peaks in control versus METTL3-knockdown condition, indicating that global translation was largely unperturbed as a result of METTL3 depletion (Figure 3A, B). Consistent with this, the distribution of the endogenous, non-methylated *hprt1* mRNA (internal control) remained constant across fractions regardless of METTL3 status (Figure 3C, lower panel). These findings indicate that global depletion of m^6^A modification does not grossly alter translation efficiency.

**Figure 3.**
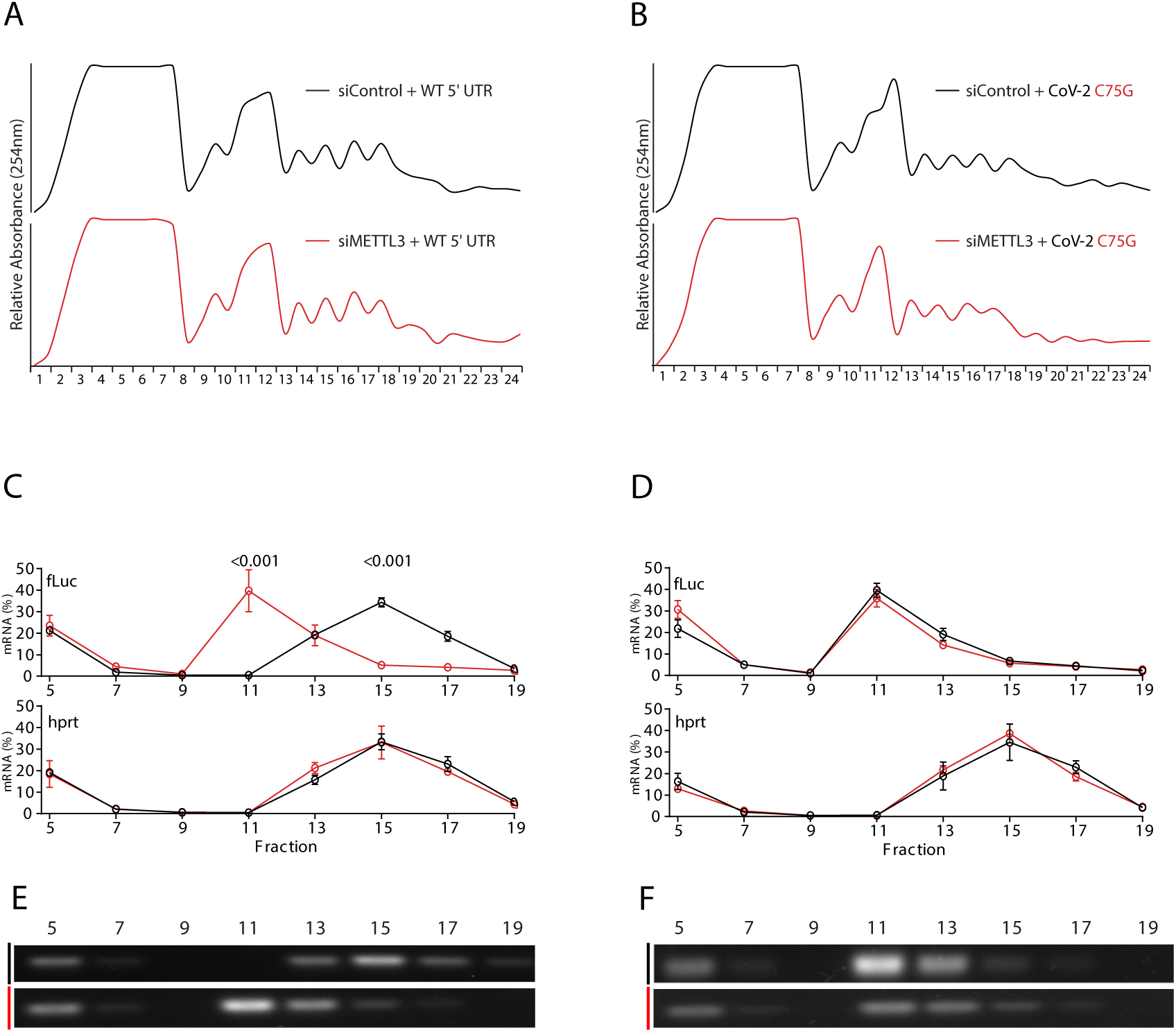
m^6^A Inhibition Impairs SARS-CoV-2 5’-UTR Promoted Translation Initiation. Ribosome profiling of HEK293T cells transfected with **A.** pGL3-CoV-2-5’-UTR (SL1-TSS)-fLuc or **B.** pGL3-CoV-2-5’-UTR (SL1-TSS)^C75G^-fLuc with METTL3 siRNA knockdown. Relative distribution (%mRNA) of fLuc (top) and hprt (bottom) determined by RT-qPCR in fractions 5-19 from **C.** pGL3-CoV-2-5’-UTR (SL1-TSS)-fLuc **D.** pGL3-CoV-2-5’-UTR^C75G^ (SL1-TSS)-fLuc transfected HEK293T cells. Results from three experiments (3 biological replicates each) shown as mean±SEM. Statistical testing performed using two-way ANOVA with Šídák’s multiple comparisons test. Representative semi-quantitative low cycle number PCR for fLuc from **E.** pGL3-CoV-2-5’-UTR (SL1-TSS)-fLuc **F.** pGL3-CoV-2-5’-UTR^C75G^ (SL1-TSS)-fLuc transfected HEK293T cells.

In stark contrast, when we examined the SARS-CoV-2 5′-UTR reporter transcripts, we observed a pronounced redistribution of ribosome occupancy toward lighter gradient fractions upon METTL3 knockdown (Figure 3C, upper panel and Figure 3E). In control cells, the viral-UTR–driven mRNA predominantly co-sedimented with heavier polysome-associated fractions, reflecting efficient translation. On the other hand, METTL3 depletion shifted the peak toward 40S/60S-associated fractions, indicating a strong inhibition of translation (Figure 3C and E). We next asked whether removal of the single m**^6^**A site at A74 was sufficient to phenocopy METTL3 depletion. Indeed, the C75G mutant reporter displayed a virtually identical shift toward lighter fractions, both in the presence and absence of METTL3 knockdown (Figure 3D, upper panel and Figure 3F), suggesting that no additional methylation sites contribute to this effect. These results demonstrate that methylation of this site is a critical determinant of translational regulation exerted by SARS-CoV-2 5′-UTR.

### m^6^A methylation promotes accessibility of SARS-COV-2 5′-UTR

Previous work has demonstrated that m^6^A modification can interfere with RNA secondary structure and destabilize local structures^55^. This prompted us to explore whether the effect of m^6^A modification at A74 on translation is related to the effect of m^6^A on the secondary structure of the 5’-UTR. Modeling the influence of m^6^A on RNA folding^60^ predicted that methylation at A74 energetically destabilizes the SARS-CoV-2 5′-UTR, particularly by disfavoring the closed conformation of stem–loop 3, SL3, (ΔG = 2.1 kcal/mol vs. 1.6 kcal/mol when methylated), while concomitantly stabilizing the open conformation (Supplementary Figure 3A, B). Consistent with this prediction, Becker et al. (2024) have reported that m^6^A methylation at A74 in the SARS-CoV-2 5′-UTR destabilizes SL3, thereby favoring the formation of TRS L:TRS B duplexes and promoting the discontinuous transcription of subgenomic mRNAs ^49^. These findings together suggest that changes in SL3 stability could influence both local stability at A74 and interactions between SL3 and other structural elements in the 5’UTR. We, therefore, set out to examine the role of structural domains in the 5’UTR by engineering a series of truncated UTR variants: one spanning stem–loops 1 through 3 (SL1–3), another including SL1 through SL4.5 (SL1–4.5), and compared them to the longer constructs ending at the transcription start site (SL1–TSS) or the fifth stem–loop (SL1–5) (Figure 4A). Each fragment was cloned upstream of the luciferase reporter, and translation was assayed in HEK293T cells under conditions that either preserve or abrogate m^6^A (via METTL3 siRNA or the A74T point mutation). All of the constructs generated robust and consistent mRNA once transfected and qPCR did not reveal any alteration in mRNA levels as a result of METTL3 knockdown (Supplementary Figure 2F).

**Figure 4.**
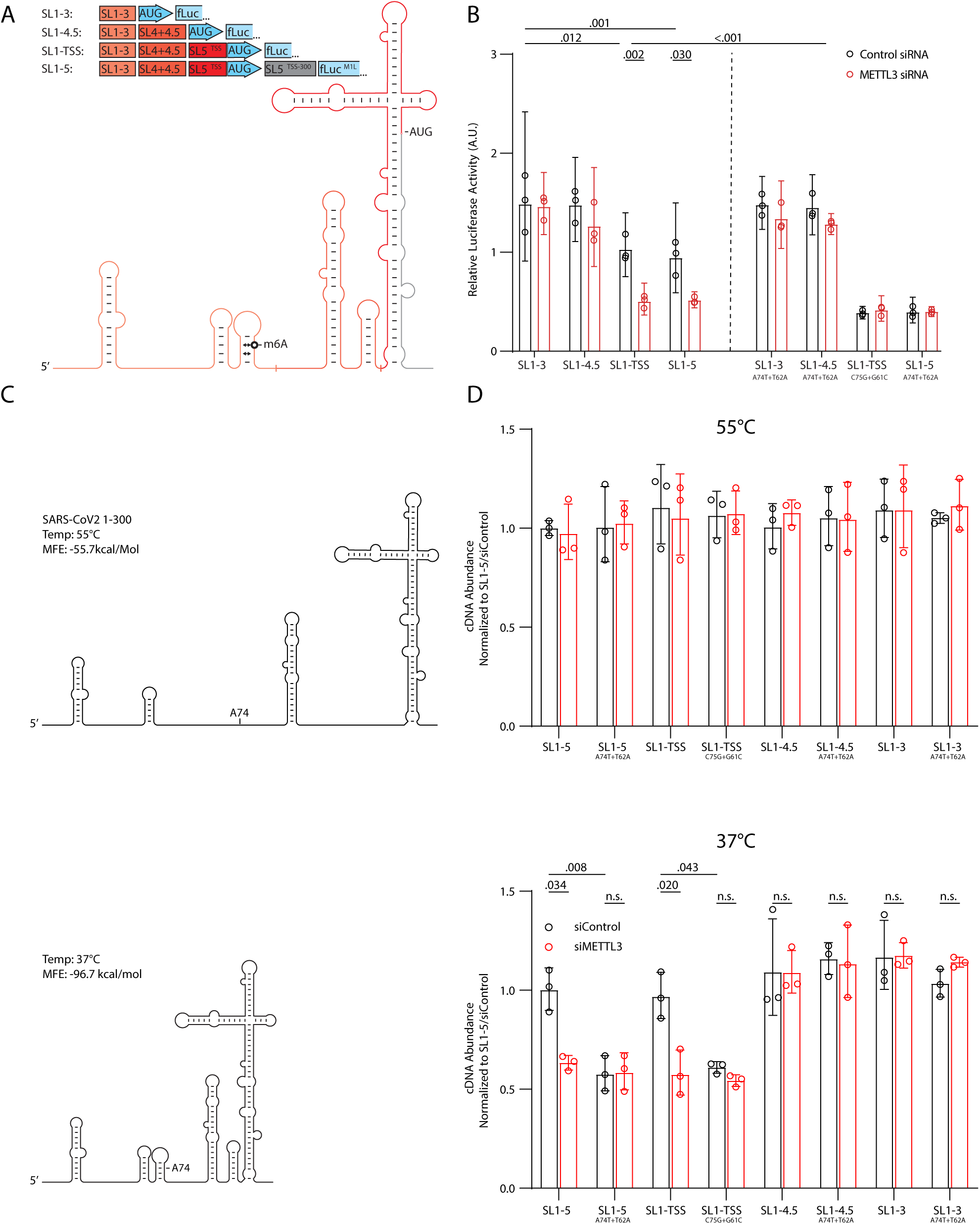
m^6^A Methylation Promotes SARS-CoV-2 5’-UTR Accessibility. **A.** Schematic of SARS-CoV-2-5’UTR segments used in this study. **B.** Luciferase assay of pGL3-CoV-2-5’-UTR (SL1-3)-fLuc, pGL3-CoV-2-5’-UTR (SL1-4.5)-fLuc, pGL3-CoV-2-5’-UTR (SL1-TSS)-fLuc, pGL3-CoV-2-5’-UTR (SL1-5)-fLuc, with or without METTL3 siRNA knockdown or mutation at A74T+T62A or C75G+G61C. Results from three experiments (3 biological replicates each) shown as geometric mean; error bars are geometric SD. Statistical testing was performed using Tukey’s HSD under two-way ANOVA. **C.** Calculated minimum free energy structure for SARS-CoV2 (SL1-5) RNA under native (37°C) or denaturing (55°C) conditions. **D.** RT-qPCR of CoV-2-5’-UTR (SL1-3)-fLuc, CoV-2-5’-UTR (SL1-4.5)-fLuc, CoV-2-5’-UTR (SL1-TSS)-fLuc, CoV-2-5’-UTR (SL1-5)-fLuc cDNA generated under denaturing (55 °C, top) and native (37 °C, bottom) with or without METTL3 siRNA knockdown or m^6^A mutation, normalized to hprt mRNA level. Results of three experiments (3 biological replicates each) normalized to CoV-2-5’-UTR (SL1-5)-fLuc and shown as geometric mean; error bars are geometric SD. Statistical testing was performed using Tukey’s HSD under two-way ANOVA.

Neither of the truncated SL1–3 and SL1–4.5 reporters exhibited any dependence on m^6^A: luciferase output remained unchanged regardless of METTL3 knockdown or A74T disruption (Figure 4B). In stark contrast, both the full-length (SL1–265) and extended (SL1–5) constructs recapitulated the robust m^6^A-dependent enhancement of translation, suggesting that the interplay between SL3 (which harbors A74) and distal elements in SL5 is required for the methylation-driven regulatory effect (Figure 4B).

These results suggested to us that a potential interaction between SL3 and SL5, when A74 is not methylated, may exert translation inhibition by hindering protein accessibility and slowing the ability of the helicase action prior to the start of translation. As an alternative to SHAPE (Selective 2′-Hydroxyl Acylation analyzed by Primer Extension)^61^, we used an RT-qPCR approach to estimate any potential interaction between m6A at A74 and other structural features in the SARS-CoV-2 5′-UTR that might influence RNA stability and thus act as hinderance to translation machinery. destabilizes. A similar method has recently been used to assay interference of small molecules with the tertiary structure of SARS-CoV-2^62^. Our approach takes advantage of a strand-displacement–deficient reverse transcriptase from Moloney murine leukemia virus (MMLV-RT)^63^ that is prone to pausing at folded structural elements^64^. By performing parallel RT reactions under denaturing (high-temperature) versus non-denaturing (low-temperature) conditions, template accessibility can be inferred: complex folds impede enzyme progression in the native setting, but impose minimal hinderance under denaturing conditions. As such, combining this reverse transcriptase assay with a qPCR strategy, we can quantify the efficiency of the RT reaction and predict which structural features correlate with more or less stable secondary structures. Following this approach, we found that under denaturing conditions (55°C), RT efficiency on the SARS-CoV-2 5′-UTR reporter was invariant across the truncated or full-length 5′-UTR reporters, and remained so after METTL3 knockdown or DRACH motif abrogation (Figure 4C-D, top). However, in non-denaturing reactions (37°C), METTL3 knockdown resulted in reduced cDNA synthesis from the longer SL1-TSS and SL1-SL5 5′-UTR RNA templates and both the C75G and A74T mutants showed a similar reduction in cDNA synthesis (Figure 4D, bottom). No further reduction occurred when combining METTL3 depletion with the point mutations, indicating that m^6^A modification at this site was the underlying cause. Additionally, truncated SARS-CoV-2 5′-UTRs containing only SL1-3 or SL1-4.5 did not show changes in reverse transcription efficiency in response to changes in m^6^A status (Figure 4D). These findings suggest that the destabilizing effect of m^6^A at A74 in SL3 can influence accessibility of protein factors such as reverse transcriptase to the full-length leader.

To further examine the effectiveness of our approach of using reverse transcription efficiency to correlate RNA secondary structure and accessibility with m^6^A modification, we turned to HEK293T cells and screened examples of human mRNAs with a single m^6^A modification exclusively in their 5’UTR^65^. We chose two human mRNAs that fit this criterion: *ACTA2* and *COX8A*. Prediction of the minimum free energy at 37°C for these two 5’UTRs compared to that of HPRT1 (control) indicates that ACTA2 has a more complex 5’UTR compared to HPRT1, while 5’UTR of COX8A appears less complex (Supplementary Figure 3C). As expected, at 55°C all three 5’UTRs are less stable with less negative ΔGs, yet maintaining the same trend (Supplementary Figure 3C). Consistently, we found that the efficiency of the RT reaction correlated with the complexity of 5’UTR for each respective gene and increasing temperature increased the success of the reaction (Supplementary Figure 3D). Strikingly, we found that under denaturing temperatures, all three mRNAs allowed for efficient reverse transcriptase activity regardless of the presence or absence of METTL3 activity; however, at 37°C knockdown of METTL3 significantly reduced the efficiency of the reverse transcriptase for both COX8A and ACTA2 but was unchanged for HPRT1 (Supplementary Figure 3C). These results further support our proposed framework for understanding the relationship between 5’UTR m^6^A modification and translational regulation and suggest that a similar process likely governs this relationship in humans.

## DISCUSSION

We demonstrate that despite its predicted complex secondary structure, the SARS-CoV-2 5′-UTR is an efficient leader sequence. A single m^6^A modification at A74 accounts for this anomaly, as its mutation or inhibition of METTL3 significantly impair the ability of SARS-CoV-2 5′-UTR to drive translation. Our experiments indicate that this m^6^A modification is particularly relevant to the efficiency of the full-length leader, responsible for facilitating translation of the polycistronic ORFs 1a and 1b when the highly complex stem loops SL5 is present. As such, the translational efficiency of the truncated versions of the leader, in the absence of SL5, do not show sensitivity to A74 mutation or inhibition of METTL3. Our work provides a mechanistic framework for understanding the epitranscriptomic control of coronavirus replication and highlights the viral 5′-UTR as a potential target for next-generation antiviral strategies against COVID-19^49^.

Recent studies have highlighted the pervasive role of m^6^A methylation in regulating the life cycle of diverse RNA viruses. On the one hand, in the case of enterovirus 71 (EV71), influenza A virus (IAV), human immunodeficiency virus (HIV), hepatitis B virus (HBV), and simian virus 40 (SV40) m^6^A methylation generally promotes replication and infectivity^40,41,66,67^. On the other, m^6^A appears to exert inhibitory effects^66^ on other viruses such as hepatitis C virus (HCV) and Zika virus (ZIKV)^40,66,68^. In the case of SARS-CoV-2, early reports yielded conflicting insights into the net impact of m^6^A: some groups observed that m^6^A deposition on viral RNA restricts replication and that depletion of METTL3, its cofactors, or YTH-domain reader proteins exacerbates infection^48^; by contrast, other work found that pharmacological blockade of m^6^A erasers (ALKBH5, FTO) suppresses viral load, whereas METTL3 knockdown or inhibition reduces infectivity^47,69^. Additional clinical and genetic studies have linked variation in m^6^A machinery components to COVID-19 disease severity and risk, further underscoring the complexity of host–virus epitranscriptomic interactions^70–75^. Our findings reconcile these seemingly discordant observations by pinpointing an essential, METTL3-dependent m^6^A mark at A74 within the viral 5′-UTR that specifically enhances translation initiation. Unlike bulk viral replication assays, our targeted approach—combining precise mutagenesis of the DRACH motif, ribosome-profiling, and RNA-structure probing—reveals that loss of A74 methylation uniformly impairs reporter-based translation without altering global host translation. Furthermore, our findings indicate an unexpected relationship between YTH cytoplasmic m^6^A readers and METTL3. While we find m^6^A modification enhances translation, the same site is used by the readers to dampen it. This suggests that while the readers require the presence of m^6^A moiety at A74 to exert their inhibitory action, methylation of A74 by METTL3 independently of YTH readers is sufficient for enhancing translation initiation, This reader-independent mechanism offers a molecular explanation for why different experimental systems, cell types, or timing of m^6^A perturbation may yield divergent outcomes: the balance between pro-viral effects of site-specific m^6^A modification and anti-viral m^6^A reader-mediated actions likely varies depending on cellular context, the ensemble of methylated sites across host and viral transcripts, and the interplay of writers, readers, and erasers in each setting.

Despite the growing catalog of m^6^A maps in SARS-CoV-2 genomes^47,48,76^ the A74 site within the TRS-L has been detected only rarely^76,77^. This underrepresentation likely stems from technical biases—such as 3′ end capture preference in sequencing workflows, platform-specific limitations of Nanopore and PacBio sequencing, and the formidable secondary-structure complexity of the 5′-UTR that impedes reverse-transcription and library preparation^78^. Moreover, A74 lies adjacent to the cap structure, raising the possibility that internal m^6^A could be misclassified as cap-proximal m^6^Am (cap-adjacent adenosine dimethylation)^79,80^. Finally, the field’s emphasis on 3′-UTR–enriched m^6^A signals in host mRNAs has historically overshadowed functional investigation of 5′-end methylation^37^. In this work, by focusing exclusively on the viral 5′-UTR and employing complementary biochemical and genetic assays, we provide definitive evidence that A74 methylation, at a site universally maintained among SARS-CoV-2 variants (Supplementary Figure 1A), confers a crucial translational advantage.

How does m^6^A at A74 promote translation? Previous work has suggested that the destabilizing effect of A74 m^6^A on SL3 may influence interaction between remote regions of mRNA^49^. Consistently, we find that A74 mutation curtails the ability of a strand-displacement–deficient reverse transcriptase to efficiently access the full-length leader sequence, suggesting that in the absence of m6A SL3 and SL5 may interact to increase the complexity of the secondary structure. From this a model emerges that a potential interaction between SL3 and SL5 likely underlies the differences we observe in translation-reporter activity. In the absence of m^6^A, the viral leader likely adopts a more rigid conformation that sterically hinders pre-initiation complex progression, thereby diminishing translation initiation. Our findings are consistent with this model: in the presence of SL5, m^6^A at A74 is critical for translation efficiency; however, when SL5 is removed, m^6^A does not influence the efficiency of translation. At the same time, mere presence of SL5 does not inhibit translation when SL3 is destabilized by m^6^A modification at A74. By linking m^6^A modification to secondary-structure dynamics and translational regulation, this work extends our foundational knowledge of m^6^A biology and points to a potential new therapeutic design to tackle SARS-CoV-2.

## MATERIALS AND METHODS

### Genomic Sequences

The complete genome sequence of the SARS-CoV-2 viral genome was obtained from NCBI’s Genbank reference sequence NC_045512. The 5′-UTR region (1-265 nt) was selected based on Genbank annotation. The complete mRNA sequence of Homo sapiens hemoglobin subunit beta (*Hbb*) was obtained from Genbank reference sequence NM_000518. The 5′-UTR Region (1-50 nt) was selected based on Genbank annotation. Genomic sequences for human 5′-UTRs were obtained from UCSC Genome Browser database, assembly GRCh37/hg19 for m^6^A annotation, and GRCh38/hg38 otherwise. 5′-UTRs were selected using the extractor functions from R package GenomicFeatures 1.48.1.

### m^6^A Site Prediction and Annotation

m^6^A sites in the SARS-CoV-2 and *Hbb* 5′-UTRs were predicted by BLAST search for DRACH Motifs (i.e. [AGU][AG]AC[ACU]), and confirmed using SRAMP^81^.

### RNA Secondary Structure and Minimum Free Energy Prediction

RNA secondary structure and minimum free energy was predicted using RNAFold 2.4.18 with default settings^82^. For structure prediction at denaturing temperature (Figure 4C), RNAFold 2.4.18 was used with temperature parameter set at 55°C. For m^6^A-dependent structure prediction, RNAStructure 6.4 with the following options: --alphabet m^6^A -m 100 -p 100^60^. R2R and Illustrator were used to visualize the predicted structures. Predicted MFE was length normalized using Trotta’s formula to compute adjusted MFE^53^:

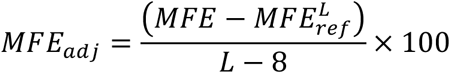

Where L is the length of the 5′-UTR and 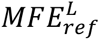 is the expected MFE for an RNA sequence of length L and GC fraction of 0.5.

### Luciferase Activity Assays

Luciferase activity assays were performed as per kit manufacturer instructions (Promega, #E1500). Briefly, cell media was aspirated and HEK293T cells washed briefly in PBS. Cells were then lysed in 1X Lysis Buffer (20 µL/well for 96-well plates or 900 µL/100mm dish). Lysate were then removed and briefly centrifuged. 20 µL of supernatant was mixed with 100 µL of Luciferase Assay Reagent and then measured on a plate reader.

### Plasmid Construction and Transfection

pGL3-HBB 5′-UTR-4X-fLuc and pGL3-SARS-CoV2-5′-UTR-fLuc plasmids were obtained as a generous gift from Ruggero Lab, UCSF. For SARS-CoV-2 5′-UTR^C75G^ and SARS-CoV-2 5′-UTR^C75G^ ^+ G61C^, reporter plasmids, gBlocks™ DNA fragments (Integrated DNA Technologies) containing 1-265 bp of SARS-CoV-2 genome with indicated mutations were flanked by restriction sites and inserted immediately upstream of fLuc into pGL3-fLuc plasmids (Promega, #E1751) using standard restriction cloning. For construction of complete and truncated reporter constructs, SARS-CoV-2 5′-UTR 1-300 bp WT and A74T or A74T+T61A mutants joined with a 6xGly flexible linker^83^ were obtained as gBlocks™ DNA fragments from which SARS-CoV-2 5′-UTR SL1-3 (1-82 bp), SL1-4.5 (1-147 bp) constructs were amplified and flanked by restriction through PCR, and then inserted in pGL3-fLuc vector using standard restriction cloning. For the SL1-5 constructs only, the initiating methionine of luciferase was mutated to lysine to ensure that only the CoV-2 TSS initiated ORF was present. Cells were transfected as per manufacturer protocol (Polyplus, #101000046). Briefly, HEK293T cells were seeded at ∼3x10^5^ cells/cm^2^ and transfected with ∼300pM plasmid DNA (0.1µg/well in 96-Well plate or 10ug/100mm dish), and 25nM siRNA (Dharmacon, #L-018095-02, #L-021009-02, and #D-001810-10) or treated with STM2457 in DMSO (Selleckchem, # S9870) after 24 hours. Gene expression and Luciferase activity were assayed 24 and 48 hours respectively after transfection.

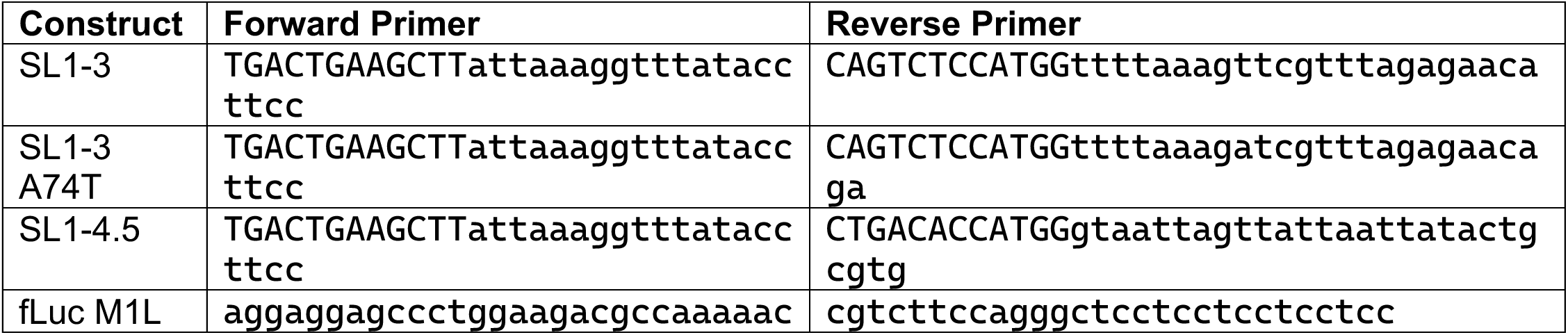

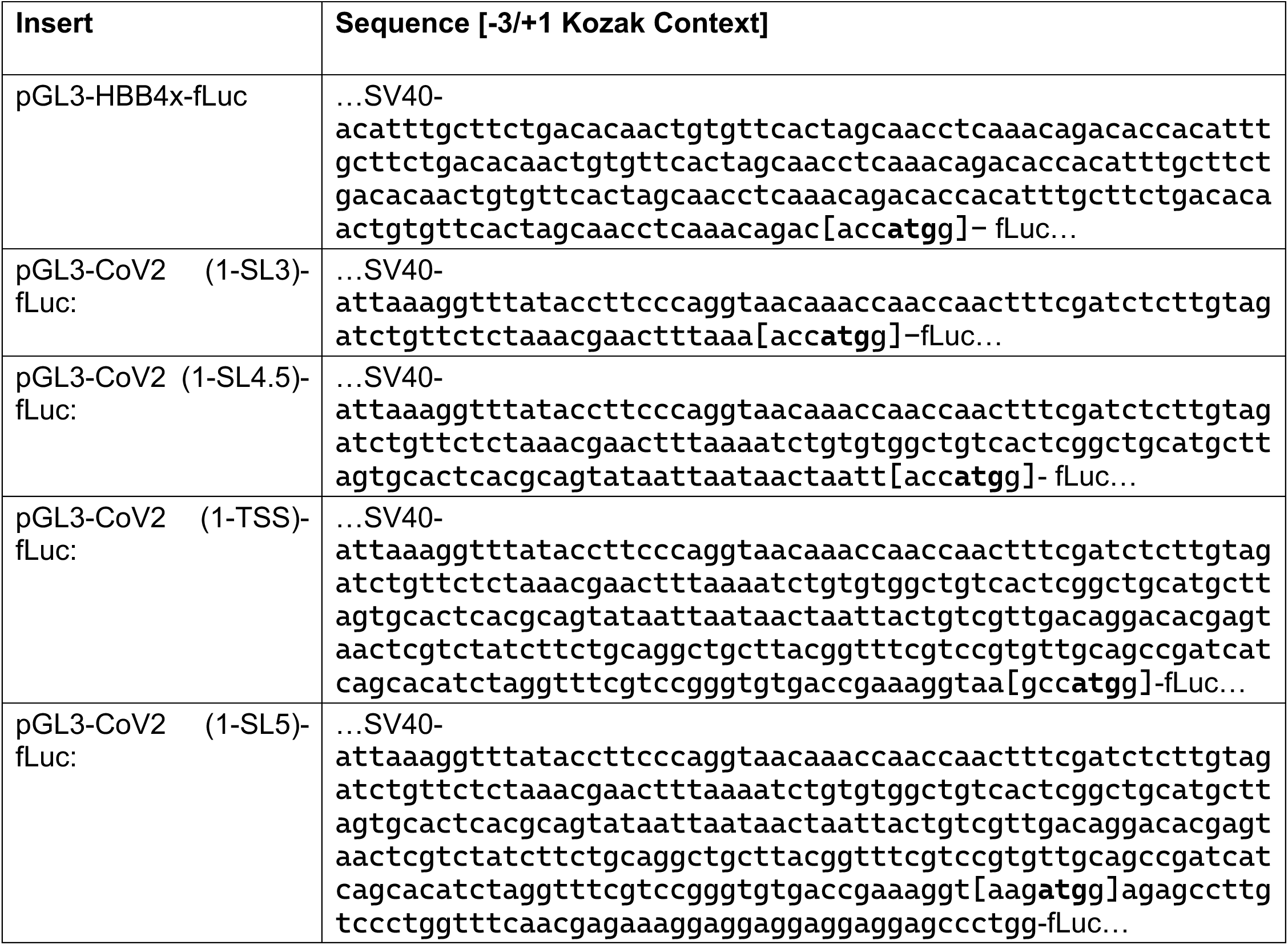

### RNA Immuno-precipitation

m^6^A RNA-IP enrichment was performed using kit manufacturer protocol (New England BioLabs, #E1610S). Briefly, Total RNA from HEK293T cells was extracted using TRIzol and fragmented using NEBNext® Magnesium RNA Fragmentation Module (New England BioLabs, E6150S). 250µg of fragmented RNA was incubated with 250 µL of m^6^A Ab conjugated to protein G magnetic beads Reaction Buffer for 1 hour at 4°C. Samples were then briefly centrifuged, supernatant was discarded, and RNA was eluted in 150 µL of clean up binding buffer and purified for downstream RT-qPCR.

### Reverse Transcription and Quantitative PCR

For cDNA production for standard qPCR, 1 µg of purified RNA was reverse transcribed using iScript cDNA synthesis Kit (BioRad, #1708891) as per manufacturer instructions where first strand cDNA synthesis was primed with a mix of poly-dT and random hexamers. For secondary structure sensitive reverse transcription reactions (Figure 4D, Supplementary Figure 3D), the same kit was used with the following modifications: the 70°C 10-minute denaturing incubation was omitted, reaction mix was supplemented with MgCl to a final concentration of 5 mM to ameliorate reduced binding of RT primers at higher temperatures, and first strand synthesis was primed with oligo(dT)_20_ primers at a final concentration of 1 µM (IDT, #51-01-15-01).

Quantitative PCR was performed using GoTaq qPCR Master Mix (Promega, #A6001) with primers:

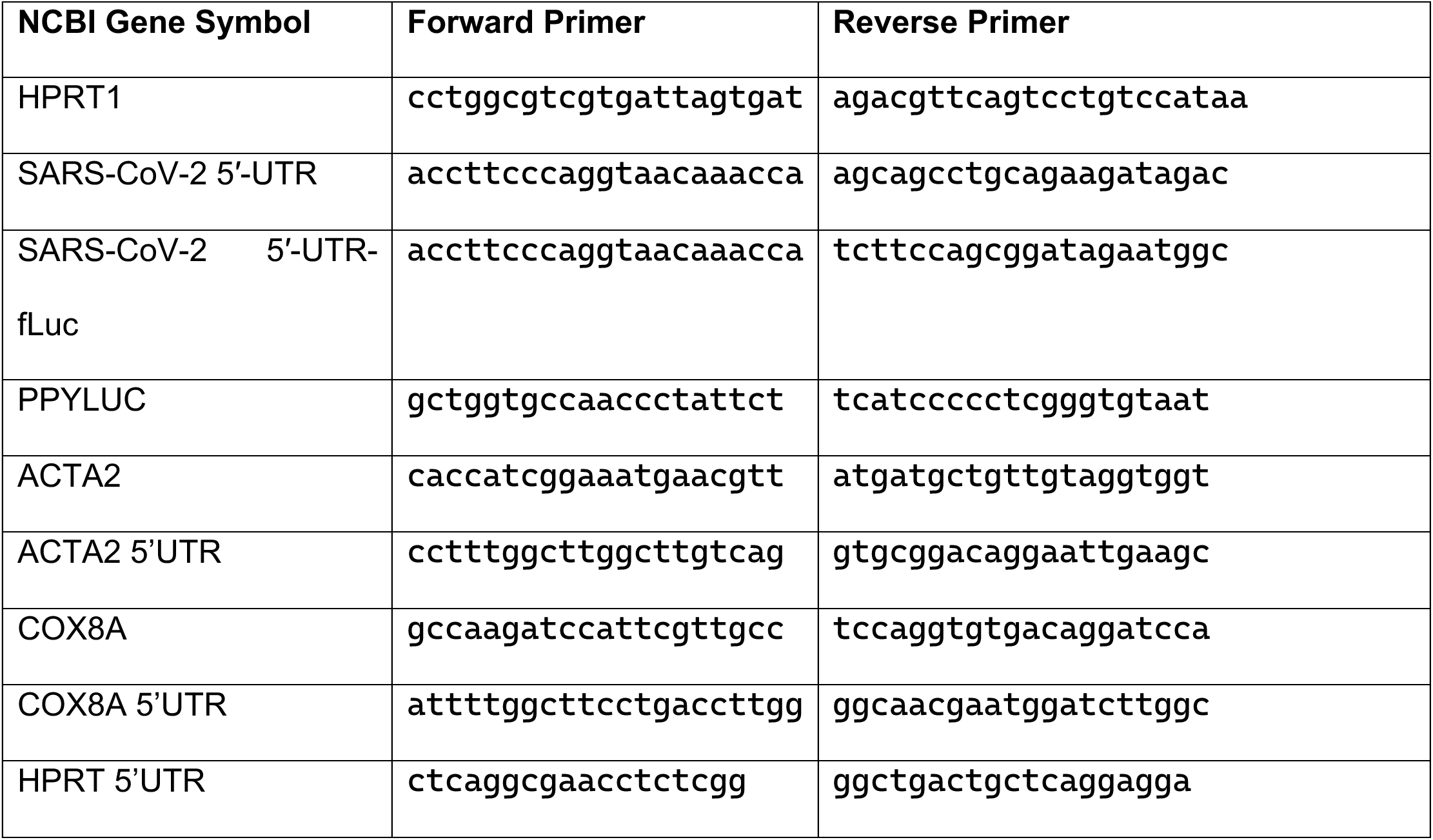

### Ribosome Profiling

HEK293T cells were seeded at ∼3x10^5^ cells/cm^2^ in a 100 mm dish and transfected as previously described. After 48 hours, Cycloheximide was added to culture media to a concentration of 50 µg/mL. After a brief 20-minute incubation, cells were scraped, pelleted, and ∼10^7^ cells were resuspended in a lysis buffer of 20 mM Tris, 10 mM MgCl_2_, 250 mM NaCl, 100 µg/mL Cycloheximide, and 0.4% NP-40 supplemented with protease inhibitor (Thermofisher, #A32963). Cells were sheared in a Dounce homogenizer, and the resultant lysate was centrifuged briefly to remove debris. Absorbance at 260nm was then measured to estimate RNA concentration in supernatant. 500 µL of 25 OD_260_ cleared lysate was carefully layered atop 20-60% sucrose gradient buffered in lysis buffer. Finally, gradients were centrifuged at 180,000 x g for 110 minutes at 4°C and subsequently fractionated.

### SARS-CoV-2 5′-UTR Mutation Analysis

Recurrent mutations track in SARS-CoV-2 were obtained from Nextstrain (accessed August 4, 2022) and viewed using UCSC Genome browser track hub^84^. Consensus sequences for SARS-CoV-2 5′-UTRs were derived from the NIH SARS-CoV-2 datahub. Briefly, all Complete Nucleotide SARS-CoV-2 sequences from obtained and grouped by WHO lineage classification, based on Pango lineage metadata. Nucleotides 1-300 from each sequence were then aligned to the SARS-CoV-2 reference 5′-UTR (NC_045512) using MAFFT 7.490 using options --6merpair --maxambiguous 0.05 –addfragments –keeplength^85^. Consensus sequences for each multiple alignment (excluding gaps) were then determined using R Biostrings 2.64.0 package^86^. Consensus secondary structure was determined using RNAFold 2.4.18, and structure diagrams were prepared using R2R. Multiple alignment figures were prepared using ESPript 3.0^87^.

## ACKNOWLEDGEMENTS

We thank the Ruggero laboratory (UCSF) for kindly providing the pGL3*-hbb-fLuc* plasmids; M. Mori for help with the polysome preparations; and M. Moujahidine for administrative support. This work was supported by an NIH grant (R01NS082793) to APH.

## AUTHOR CONTRIBUTIONS

A.A. and A.P.H. designed the experiments. A.A., M.C., and G.S. performed the experiments. A.A. and

A.P.H. prepared the manuscript. A.A., G.S. and A.P.H. edited the manuscript.

## COMPETING INTERESTS

The authors declare no competing interests.

## MATERIALS AND CORRESPONDENCE

Correspondence and requests for materials should be addressed to Pejmun Haghighi.

---

**Supplementary Figure 1.**
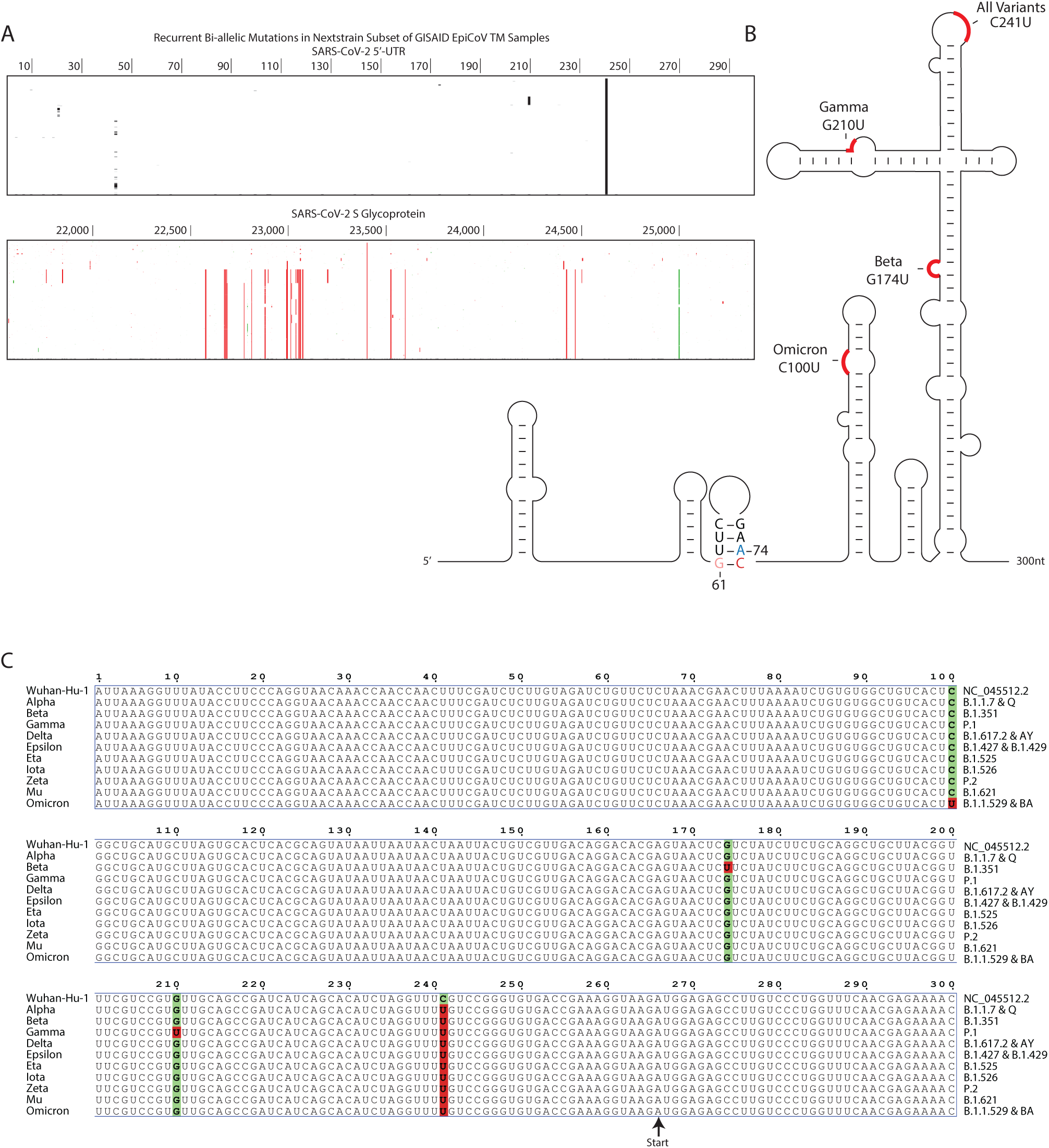
SARS-CoV-2 5’-UTR is Highly Conserved Among SARS-CoV-2 Variants. **A.** Genome browser tracks showing Nextstrain recurrent mutations in SARS-CoV-2 5’-UTR (top) and SARS-CoV-2 S Glycoprotein (bottom). Non-synonymous mutations in protein coding regions are shown in red, and synonymous mutations are shown in green. **B.** Lineage-specific SARS-CoV-2 5-’UTR mutations for all WHO variants of interest and variants of concern mapped to predicted secondary structure. **C.** Lineage-specific multiple alignment of consensus sequences to reference SARS-CoV-2 genome (Wuhan-Hu-1; NC_045512.2) for all WHO variants of interest and variants of concern. Polymorphism sites are highlighted in green, with mutant allele highlighted in red.

**Supplementary Figure 2.**
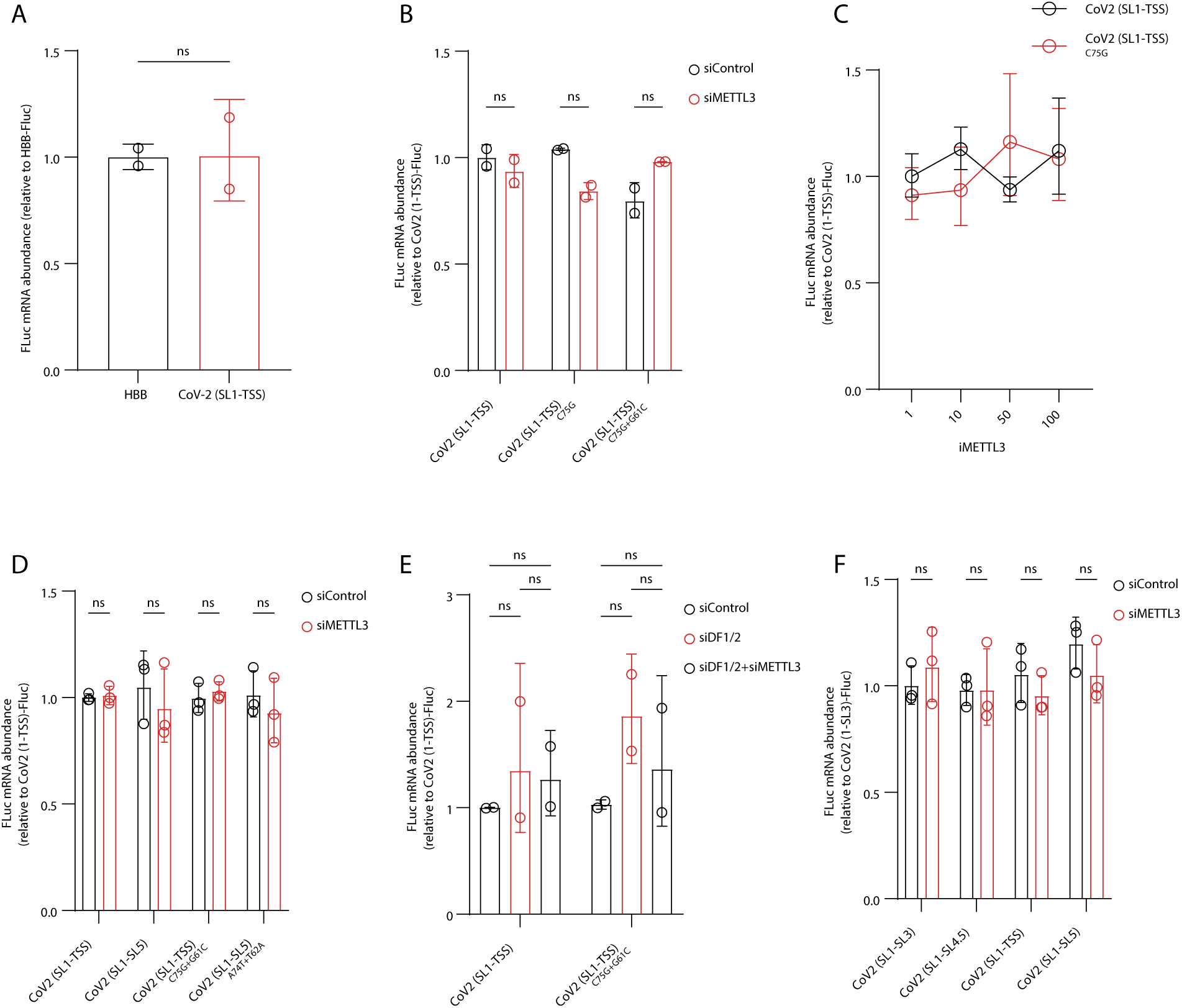
m^6^A Methylation Does not Alter mRNA levels of pGL3-SARS-CoV-2 5’UTR-fLuc. **A.** qPCR measurement of HBB-fLuc and CoV-2 (SL1-TSS)-fLuc mRNA in transfected HEK293T cells. Results from 2 experiments (3 biological replicates each) shown as geometric mean; error bars are geometric SD. Statistical testing performed using Welch’s T-Test. **B.** qPCR measurement of CoV-2 (SL1-TSS)-fLuc mRNA with or without METTL3 siRNA or C75G/G61C mutations in transfected HEK293T cells. Results from 2 experiments (3 biological replicates each) shown as geometric mean; error bars are geometric SD. Statistical testing performed using two-way ANOVA with Šidák’s multiple comparison test. **C.** qPCR measurement of CoV-2 (SL1-TSS)-fLuc mRNA with or without C75G mutations in HEK293T cells treated with STM2457. Results from a single experiment (3 biological replicates) shown. **D.** qPCR measurement of CoV-2 (SL1-TSS)-fLuc or CoV-2 (SL1-5)-fLuc with or without METTL3 siRNA or C75G+G61C or A74T+T62A mutation as indicated in transfected HEK293T cells. Results from 3 experiments (3 biological replicates each) shown as geometric mean; error bars are geometric SD. Statistical testing performed using two-way ANOVA with Tukey’s HSD. **E.** qPCR measurement of CoV-2 (SL1-TSS)-fLuc with or without METTL3 and DF1/2 siRNA or C75G+G61C or A74T+T62A mutation as indicated in transfected HEK293T Cells. Results from 2 experiments (3 biological replicates each) shown as geometric mean; error bars are geometric SD. Statistical testing performed using two-way ANOVA with Tukey’s HSD. **F.** qPCR measurement of CoV-2 (SL1-3)-fLuc, CoV-2 (SL1-4.5)-fLuc, CoV-2 (SL1-TSS)-fLuc, or CoV-2 (SL1-5)-fLuc with or without METTL3 siRNA in transfected HEK293T cells. Results from 3 experiments (3 biological replicates each) shown as geometric mean; error bars are geometric SD. Statistical testing performed using two-way ANOVA with Šidák’s multiple comparison test. n.s. indicates p-value > 0.05. For all panels hprt was used as internal normalization control.

**Supplementary Figure 3.**
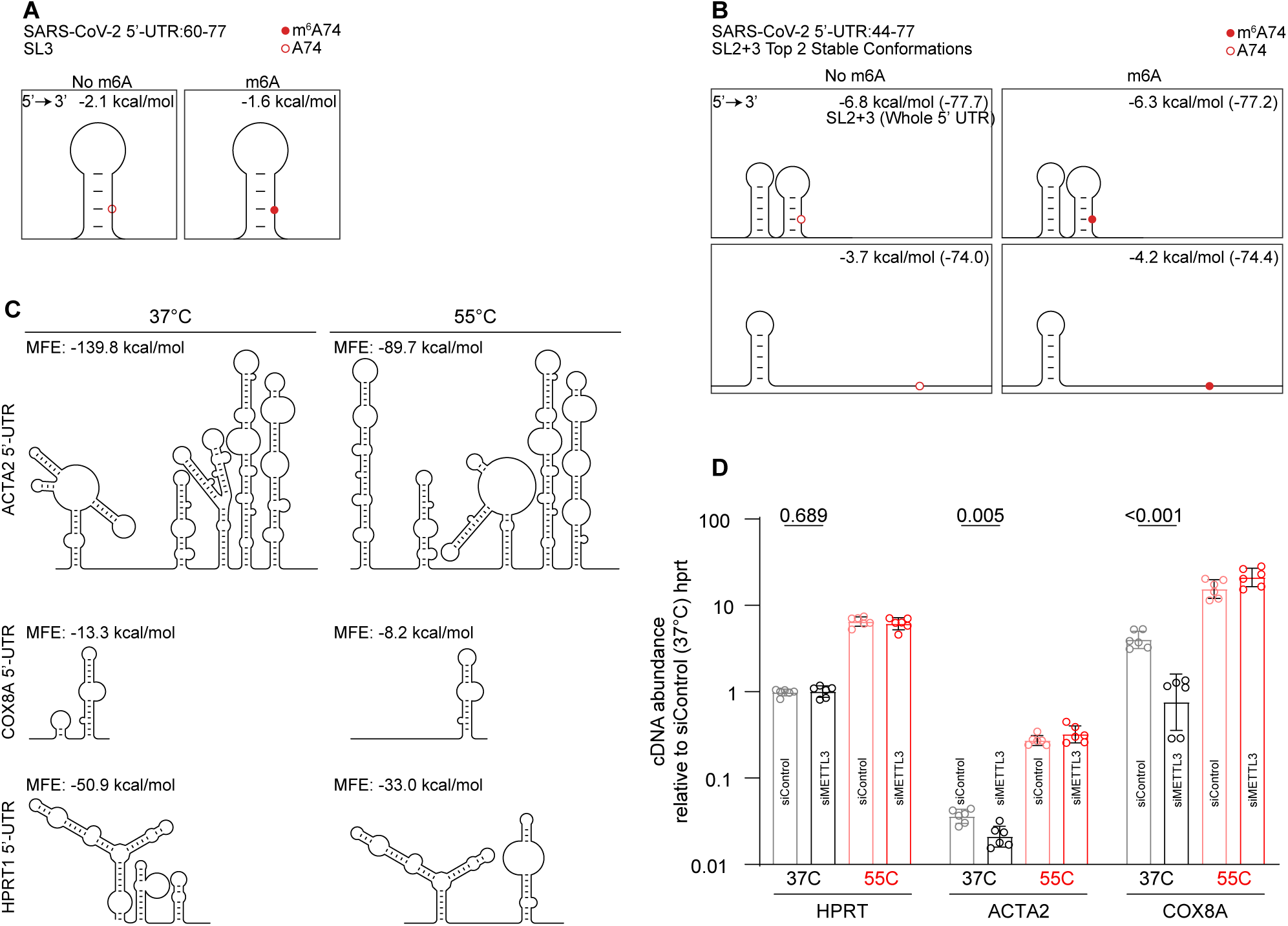
m^6^A Methylation Destabilizes SARS-CoV-2 5’-UTR Secondary Structure. **A.** m^6^A-dependent minimum free energy secondary structure prediction of stem loop 3 using RNAstructure with m^6^A alphabet. **B.** m^6^A-dependent minimum free energy secondary structure prediction of stem loops 2+3 using RNAstructure with m^6^A alphabet. **C.** Minimum free energy secondary structure prediction of *acta2, cox8a,* and *hprt* 5’-UTRs at 37°C and 55°C. **D.** cDNA abundance of *acta2, cox8a,* and *hprt* mRNA from HEK293T cells with or without METTL3 siRNA knockdown reverse transcribed under native (37°C) or denaturing (55°C) conditions normalized to *hprt*. Multiple t-test with Welch correction, and multiple testing adjustment with Holm-Šídák method

## Notes

### Competing Interest Statement

The authors have declared no competing interest.

### Summary of Updates

The manuscript has been updated to reflect new developments in the literature on SARS-CoV-2 biology and includes new strengthening experiments. Particularly, Figures 2, 4, and supplementary figure 3 contain new results. A new supplementary figure 2 contains results from additional control experiments. All text and figures have been edited for clarity.

## REFERENCES

1. Ciotti, M., Ciccozzi, M., Pieri, M. & Bernardini, S. The COVID-19 pandemic: viral variants and vaccine efficacy. Critical Reviews in Clinical Laboratory Sciences 59, 66–75 (2022).

2. Shinde, V. et al. Efficacy of NVX-CoV2373 Covid-19 Vaccine against the B.1.351 Variant. N Engl J Med **384**, 1899–1909 (2021).

3. Rubin, R. COVID-19 Vaccines vs Variants-Determining How Much Immunity Is Enough. JAMA 325, 1241–1243 (2021).

4. Haque, A. & Pant, A. B. Mitigating Covid-19 in the face of emerging virus variants, breakthrough infections and vaccine hesitancy. Journal of Autoimmunity 127, 102792 (2022).

5. Zhang, X. et al. A spatial vaccination strategy to reduce the risk of vaccine-resistant variants. PLoS computational biology 18, e1010391 (2022).

6. de Souza, A. S. et al. Severe Acute Respiratory Syndrome Coronavirus 2 Variants of Concern: A Perspective for Emerging More Transmissible and Vaccine-Resistant Strains. Viruses 14, 827 (2022).

7. Wang, R., Chen, J. & Wei, G.-W. Mechanisms of SARS-CoV-2 evolution revealing vaccine-resistant mutations in Europe and America. The journal of physical chemistry letters 12, 11850–11857 (2021).

8. Kim, D. et al. The Architecture of SARS-CoV-2 Transcriptome. Cell 181, 914–921.e10 (2020).

9. Wang, D. et al. The SARS-CoV-2 subgenome landscape and its novel regulatory features. Molecular Cell 81, 2135–2147.e5 (2021).

10. WHO COVID-19 dashboard. WHO COVID-19 Dashboard https://data.who.int/dashboards/covid19/.

11. Li, P., et al. Neutralization escape, infectivity, and membrane fusion of JN.1-derived SARS-CoV-2 SLip, FLiRT, and KP.2 variants. Cell Reports 43, (2024).

12. Nasir, A. et al. Predictive Modeling of Immune Escape and Antigenic Grouping of SARS-CoV-2 Variants. 2025.05.28.656328 Preprint at 10.1101/2025.05.28.656328 (2025).

13. Magazine, N. et al. Mutations and evolution of the SARS-CoV-2 spike protein. Viruses 14, 640 (2022).

14. Rosenbloom, D. I. S., Hill, A. L., Rabi, S. A., Siliciano, R. F. & Nowak, M. A. Antiretroviral dynamics determines HIV evolution and predicts therapy outcome. Nat Med 18, 1378–1385 (2012).

15. Terrault, N. A. et al. Update on prevention, diagnosis, and treatment of chronic hepatitis B: AASLD 2018 hepatitis B guidance. Hepatology 67, 1560–1599 (2018).

16. Jong, R. M. et al. Mucosal Vaccination with Cyclic Dinucleotide Adjuvants Induces Effective T Cell Homing and IL-17–Dependent Protection against *Mycobacterium tuberculosis* Infection. The Journal of Immunology 208, 407–419 (2022).

17. Bozic, I. et al. Evolutionary dynamics of cancer in response to targeted combination therapy. eLife 2, e00747 (2013).

18. Sola, I., Almazán, F., Zúñiga, S. & Enjuanes, L. Continuous and Discontinuous RNA Synthesis in Coronaviruses. Annu Rev Virol 2, 265–288 (2015).

19. 19. Masters, P. S. The Molecular Biology of Coronaviruses. in Advances in Virus Research vol. 66 193–292 (Academic Press, 2006).

20. Zúñiga, S., Sola, I., Alonso, S. & Enjuanes, L. Sequence Motifs Involved in the Regulation of Discontinuous Coronavirus Subgenomic RNA Synthesis. J Virol 78, 980–994 (2004).

21. Miao, Z., Tidu, A., Eriani, G. & Martin, F. Secondary structure of the SARS-CoV-2 5’-UTR. RNA Biology 18, 447–456 (2021).

22. Sosnowski, P., Tidu, A., Eriani, G., Westhof, E. & Martin, F. Correlated sequence signatures are present within the genomic 5′UTR RNA and NSP1 protein in coronaviruses. RNA 28, 729–741 (2022).

23. Rangan, R. et al. RNA genome conservation and secondary structure in SARS-CoV-2 and SARS-related viruses: a first look. RNA 26, 937–959 (2020).

24. Babendure, J. R., Babendure, J. L., Ding, J.-H. & Tsien, R. Y. Control of mammalian translation by mRNA structure near caps. RNA 12, 851–861 (2006).

25. Sonenberg, N. & Hinnebusch, A. G. Regulation of Translation Initiation in Eukaryotes: Mechanisms and Biological Targets. Cell 136, 731–745 (2009).

26. Hinnebusch, A. G., Ivanov, I. P. & Sonenberg, N. Translational control by 5′-untranslated regions of eukaryotic mRNAs. Science 352, 1413–1416 (2016).

27. Andrews, R. J., et al. A map of the SARS-CoV-2 RNA structurome. NAR Genomics and Bioinformatics 3, (2021).

28. Kadam, S. B., Sukhramani, G. S., Bishnoi, P., Pable, A. A. & Barvkar, V. T. SARS-CoV-2, the pandemic coronavirus: Molecular and structural insights. Journal of Basic Microbiology 61, 180–202 (2021).

29. Vögele, J. et al. High-resolution structure of stem-loop 4 from the 5′-UTR of SARS-CoV-2 solved by solution state NMR. Nucleic Acids Res 51, 11318–11331 (2023).

30. Huang, T. et al. Analysis and Prediction of Translation Rate Based on Sequence and Functional Features of the mRNA. PLOS ONE 6, e16036 (2011).

31. Leppek, K., Das, R. & Barna, M. Functional 5′ UTR mRNA structures in eukaryotic translation regulation and how to find them. Nat Rev Mol Cell Biol 19, 158–174 (2018).

32. Livingstone, M., Atas, E., Meller, A. & Sonenberg, N. Mechanisms governing the control of mRNA translation. Phys. Biol. 7, 021001 (2010).

33. Rubio, C. A. et al. Transcriptome-wide characterization of the eIF4A signature highlights plasticity in translation regulation. Genome Biology 15, 476 (2014).

34. Blanco, J. D., Hernandez-Alias, X., Cianferoni, D. & Serrano, L. In silico mutagenesis of human ACE2 with S protein and translational efficiency explain SARS-CoV-2 infectivity in different species. PLOS Computational Biology 16, e1008450 (2020).

35. Kumar, P. et al. Clinically observed deletions in SARS-CoV-2 Nsp1 affect its stability and ability to inhibit translation. FEBS Letters 596, 1203–1213 (2022).

36. Mohammadi-Dehcheshmeh, M. et al. A Transcription Regulatory Sequence in the 5′ Untranslated Region of SARS-CoV-2 Is Vital for Virus Replication with an Altered Evolutionary Pattern against Human Inhibitory MicroRNAs. Cells 10, 319 (2021).

37. Meyer, K. D. & Jaffrey, S. R. Rethinking m6A Readers, Writers, and Erasers. Annu. Rev. Cell Dev. Biol. 33, 319–342 (2017).

38. Desrosiers, R. C., Friderici, K. H. & Rottman, F. M. Characterization of Novikoff hepatoma mRNA methylation and heterogeneity in the methylated 5’ terminus. Biochemistry 14, 4367–4374 (1975).

39. Kim, G.-W. & Siddiqui, A. Hepatitis B virus X protein recruits methyltransferases to affect cotranscriptional N6-methyladenosine modification of viral/host RNAs. Proceedings of the National Academy of Sciences 118, e2019455118 (2021).

40. Kim, G.-W. & Siddiqui, A. N6-methyladenosine modification of HCV RNA genome regulates cap-independent IRES-mediated translation via YTHDC2 recognition. Proc. Natl. Acad. Sci. U.S.A. 118, e2022024118 (2021).

41. Kim, G.-W., Imam, H., Khan, M. & Siddiqui, A. N6-Methyladenosine modification of hepatitis B and C viral RNAs attenuates host innate immunity via RIG-I signaling. J Biol Chem 295, 13123–13133 (2020).

42. Manners, O., Baquero-Perez, B. & Whitehouse, A. m6A: Widespread regulatory control in virus replication. Biochimica et Biophysica Acta (BBA) - Gene Regulatory Mechanisms 1862, 370–381 (2019).

43. Brocard, M., Ruggieri, A. & Locker, N. 2017. m6A RNA methylation, a new hallmark in virus-host interactions. Journal of General Virology 98, 2207–2214.

44. Gokhale, N. S. et al. N6-Methyladenosine in Flaviviridae Viral RNA Genomes Regulates Infection. Cell Host and Microbe 20, 654–665 (2016).

45. Körtel, N. et al. Deep and accurate detection of m6A RNA modifications using miCLIP2 and m6Aboost machine learning. Nucleic Acids Research 49, e92–e92 (2021).

46. Zaccara, S. & Jaffrey, S. R. A Unified Model for the Function of YTHDF Proteins in Regulating m6A-Modified mRNA. Cell 181, 1582–1595.e18 (2020).

47. Li, N. et al. METTL3 regulates viral m6A RNA modification and host cell innate immune responses during SARS-CoV-2 infection. Cell Rep 35, 109091 (2021).

48. Liu, J. et al. The m6A methylome of SARS-CoV-2 in host cells. Cell Res 31, 404–414 (2021).

49. Becker, M. A. et al. m6A Methylation of Transcription Leader Sequence of SARS-CoV-2 Impacts Discontinuous Transcription of Subgenomic mRNAs. Chemistry – A European Journal 30, e202401897 (2024).

50. Karollus, A., Avsec, Ž. & Gagneur, J. Predicting mean ribosome load for 5’UTR of any length using deep learning. PLOS Computational Biology 17, e1008982 (2021).

51. Corbett, K. S. et al. SARS-CoV-2 mRNA vaccine design enabled by prototype pathogen preparedness. Nature 586, 567–571 (2020).

52. Polack, F. P. et al. Safety and Efficacy of the BNT162b2 mRNA Covid-19 Vaccine. N Engl J Med 383, 2603–2615 (2020).

53. Trotta, E. On the Normalization of the Minimum Free Energy of RNAs by Sequence Length. PLOS ONE 9, e113380 (2014).

54. Roost, C. et al. Structure and Thermodynamics of N ^6^ -Methyladenosine in RNA: A Spring-Loaded Base Modification. J. Am. Chem. Soc. 137, 2107–2115 (2015).

55. Spitale, R. C. et al. Structural imprints in vivo decode RNA regulatory mechanisms. Nature 519, 486–490 (2015).

56. Fan, R. et al. A combined deep learning framework for mammalian m6A site prediction. Cell Genomics 4, (2024).

57. Wacker, A. et al. Secondary structure determination of conserved SARS-CoV-2 RNA elements by NMR spectroscopy. Nucleic Acids Res 48, 12415–12435 (2020).

58. Lan, T. C. T. et al. Secondary structural ensembles of the SARS-CoV-2 RNA genome in infected cells. Nat Commun 13, 1128 (2022).

59. Zhang, Y. et al. In vivo structure and dynamics of the SARS-CoV-2 RNA genome. Nat Commun 12, 5695 (2021).

60. Kierzek, E. et al. Secondary structure prediction for RNA sequences including N6-methyladenosine. Nat Commun 13, 1271 (2022).

61. Siegfried, N. A., Busan, S., Rice, G. M., Nelson, J. A. & Weeks, K. M. RNA motif discovery by SHAPE and mutational profiling (SHAPE-MaP). Nature methods 11, 959–65 (2014).

62. Zafferani, M., Muralidharan, D., Montalvan, N. I. & Hargrove, A. E. RT-qPCR as a screening platform for mutational and small molecule impacts on structural stability of RNA tertiary structures. RSC Chem Biol 3, 905–915 (2022).

63. Okano, H. et al. Enhanced detection of RNA by MMLV reverse transcriptase coupled with thermostable DNA polymerase and DNA/RNA helicase. Enzyme Microb Technol 96, 111–120 (2017).

64. Malik, O., Khamis, H., Rudnizky, S., Marx, A. & Kaplan, A. Pausing kinetics dominates strand-displacement polymerization by reverse transcriptase. Nucleic Acids Res 45, 10190–10205 (2017).

65. Tang, Y. et al. m6A-Atlas: a comprehensive knowledgebase for unraveling the N6-methyladenosine (m6A) epitranscriptome. Nucleic acids research 49, D134–D143 (2021).

66. Williams, G. D., Gokhale, N. S. & Horner, S. M. Regulation of Viral Infection by the RNA Modification N6-Methyladenosine. Annu Rev Virol 6, 235–253 (2019).

67. Hao, H. et al. N6-methyladenosine modification and METTL3 modulate enterovirus 71 replication. Nucleic Acids Res 47, 362–374 (2019).

68. Sacco, M. T., Bland, K. M. & Horner, S. M. WTAP Targets the METTL3 m6A-Methyltransferase Complex to Cytoplasmic Hepatitis C Virus RNA to Regulate Infection. J Virol 96, e0099722 (2022).

69. Burgess, H. M. et al. Targeting the m6A RNA modification pathway blocks SARS-CoV-2 and HCoV-OC43 replication. Genes Dev. 35, 1005–1019 (2021).

70. Meng, Y. et al. RBM15-mediated N6-methyladenosine modification affects COVID-19 severity by regulating the expression of multitarget genes. Cell Death Dis 12, 1–10 (2021).

71. An, S. et al. Systematic analysis of clinical relevance and molecular characterization of m6A in COVID-19 patients. Genes & Diseases 9, 1170–1173 (2022).

72. Lu, L. et al. The risk of COVID-19 can be predicted by a nomogram based on m6A-related genes. *Infection*, Genetics and Evolution 106, 105389 (2022).

73. Chen, Y., Qin, Y., Fu, Y., Gao, Z. & Deng, Y. Integrated Analysis of Bulk RNA-Seq and Single-Cell RNA-Seq Unravels the Influences of SARS-CoV-2 Infections to Cancer Patients. International Journal of Molecular Sciences 23, 15698 (2022).

74. Zhang, T. et al. N6-methyladenosine regulates RNA abundance of SARS-CoV-2. Cell Discov 7, 1–4 (2021).

75. Zhang, X. et al. Methyltransferase-like 3 Modulates Severe Acute Respiratory Syndrome Coronavirus-2 RNA N6-Methyladenosine Modification and Replication. mBio 12, e0106721 (2021).

76. Malbec, L., et al. The RNA Demethylase FTO Controls m ^6^ A Marking on SARS-CoV-2 and Classifies COVID-19 Severity in Patients. http://biorxiv.org/lookup/doi/10.1101/2022.06.27.497749 (2022) doi:10.1101/2022.06.27.497749.

77. Liang, J. et al. How the Replication and Transcription Complex Functions in Jumping Transcription of SARS-CoV-2. Frontiers in Genetics 13, (2022).

78. Shi, H. et al. Bias in RNA-seq Library Preparation: Current Challenges and Solutions. Biomed Res Int 2021, 6647597 (2021).

79. Sun, H. et al. m6Am-seq reveals the dynamic m6Am methylation in the human transcriptome. Nat Commun 12, 4778 (2021).

80. McIntyre, A. B. R. et al. Limits in the detection of m6A changes using MeRIP/m6A-seq. Sci Rep 10, 6590 (2020).

81. Zhou, Y., Zeng, P., Li, Y.-H., Zhang, Z. & Cui, Q. SRAMP: prediction of mammalian N6-methyladenosine (m6A) sites based on sequence-derived features. Nucleic Acids Research 44, e91–e91 (2016).

82. Lorenz, R. et al. ViennaRNA Package 2.0. Algorithms for Molecular Biology 6, 26 (2011).

83. Chen, X., Zaro, J. & Shen, W.-C. Fusion Protein Linkers: Property, Design and Functionality. Adv Drug Deliv Rev 65, 1357–1369 (2013).

84. Hadfield, J. et al. Nextstrain: real-time tracking of pathogen evolution. Bioinformatics 34, 4121–4123 (2018).

85. Nakamura, T., Yamada, K. D., Tomii, K. & Katoh, K. Parallelization of MAFFT for large-scale multiple sequence alignments. Bioinformatics 34, 2490–2492 (2018).

86. Pagès, H., Aboyoun, P., Gentleman, R. & DebRoy, S. Biostrings: Efficient manipulation of biological strings. Bioconductor version: Release (3.15) 10.18129/B9.bioc.Biostrings (2022).

87. Robert, X. & Gouet, P. Deciphering key features in protein structures with the new ENDscript server. Nucleic Acids Res 42, W320–324 (2014).

